# Neural Convergence of Graded Belief and Binary Belief Uncertainty

**DOI:** 10.1101/2025.07.14.664652

**Authors:** Martin Fungisai Gerchen, Samantha Ullrich, Mathis Lessau, Peter Kirsch

## Abstract

Belief was defined by William James as “the psychological process or function of cognizing reality”, and the recent philosophical literature has emphasized that there are two types of belief: categorical and graded. The relationship between these two belief types is complex, and it is often assumed that the degrees of graded belief reflect confidence. However, this claim has not yet been addressed empirically. In this functional magnetic resonance imaging study we let N=29 young healthy participants rate their graded belief in propositions from the conspiracy theory spectrum and estimated certainty scores from the ratings that we associated with brain activation during the presentation of the propositions. We found associations of uncertainty (i.e. negative associations with certainty scores) with brain activation in the dorsomedial prefrontal cortex, right dorsolateral prefrontal cortex, and posterior parietal cortex. The clusters in the dorsomedial and dorsolateral prefrontal cortex replicated prior associations with explicit ratings of uncertainty in binary belief decisions from a study on domain-general belief and the dorsomedial prefrontal cortex effect, localized in the anterior pre-supplementary motor area, corresponds to the concepts of Alexander Bain about the close relationship between belief and action and a central role of doubt in belief. Our results provide empirical evidence for a link between graded belief and uncertainty in human cognition and emphasize the role of doubt as a relevant process to take into account in concepts of graded belief.

**Highlights:**

- Converging neural associations of graded belief levels and belief uncertainty
- Conceptual replication of belief uncertainty associations with a modified procedure
- Correspondence to classical concepts of belief by Alexander Bain and William James
- Our results emphasize doubt as a relevant process in graded belief

## 1. Introduction

Belief, defined by William James as “the psychological process or function of cognizing reality” [1, p. 283], has been debated in psychology and philosophy for several centuries [2], and more recently also in cognitive neuroscience [3]. It plays an important role in a wide variety of domains from daily decisions over mental disorders to political attitudes and conspiracy theories (CT) [4-7].

In the recent philosophical literature two types of belief are contrasted: A categorical binary (true/false) type, often extended by a neutral stance as a third option, and a graded type where beliefs are ordered by strength, also called degreed doxastic attitude or partial belief (see [8-11]), and there have been intensive discussions which is more fundamental, whether each type even exists, and how they could be integrated (see [8,11-17]).

In an influential paper, Foley [9], referencing John Locke’s “An Essay Concerning Human Understanding” [18], proposed the “Lockean thesis” to link the belief types. In this concept, a rational belief is one for which the subjective rational degree of confidence is sufficiently high above some threshold, and it has become common in the philosophical literature to assume that graded belief is reflecting confidence [10,11,17]. However, as Moon [16] points out, this equivalence is an intuitive assumption made without further evidence.

It is therefore relevant to elucidate the involved processes. Empirically, both types of belief are readily measured in humans by binary (true/false) decisions or by truths rating from 0–100%. To investigate the neural implementation of the involved cognitive processes, neuroimaging approaches like functional magnetic resonance imaging (fMRI) provide important tools to identify associations with relevant (sub)processes and differentiate them.

In a seminal fMRI study, Harris et al. [19] presented propositions (short statements) and let participants judge categorically whether they considered them as true (belief), false (disbelief), or undecidable (uncertainty). They identified associations in ventromedial prefrontal cortex (vmPFC) for belief, in inferior frontal gyrus, middle frontal gyrus, and anterior insula for disbelief, and in anterior cingulate cortex and dorsomedial prefrontal cortex (dmPFC) for uncertainty.

In an adapted version of Harris’ experiment, we separated proposition presentation, categorical true/false belief decision, and certainty judgement [2]. With this procedure we replicated the vmPFC effect of Harris et al. [19], found an association in anterior temporal cortex with disbelief, and identified distinct networks linearly associated with certainty and uncertainty. The uncertainty network was centered at the dmPFC, constituting a conceptual replication of Harris’ uncertainty effect. In a subsequent study, we investigated self-referential belief and replicated the vmPFC belief and dmPFC uncertainty effects [20].

Interestingly, the dmPFC has been implicated by Kaplan et al. [21] in resistance to belief change. In this study, the higher dmPFC activation was during the presentation of challenges contradicting strongly held views of their participants, the smaller was the belief change. While Kaplan et al. [21] interpreted their finding as reflecting the role of the dmPFC in cognitive reappraisal and regulating negative affect, it would also be consistent with an underlying uncertainty effect.

To further investigate these processes, we use an adaptation of the experiment of Kaplan et al. [21] and let participants rate their graded belief in 16 CTs from 0-100% and presented either supporting or contradicting statements for each. While the overall aim of our study was to identify brain processes associated with processing of CTs, here we focus solely on an independent pregregistered secondary analysis to identify brain activation associated with belief uncertainty.

To compare uncertainty effects estimated from graded belief to uncertainty in binary belief decisions, we present our results in conjunction with those reported in Gerchen et al. [2].

## 2. Methods

### 2.1 Sample

A group of 30 healthy volunteers participated in the study. Exclusion criteria were no MRI eligibility, neurological or acute psychiatric illness, a history of substance dependence other than nicotine, left-handedness and insufficient German language skills. One person was excluded from the analysis because of excessive head motion during fMRI scanning. Thus, the final sample size was N=29 participants (mean age 24.97 years (SD = 4.48, range 19-36 years), 22 female). 19 participants were students, 7 had an academic degree and 3 had a completed apprenticeship. Before participants were included, a screening telephone interview was conducted. Following successful screening, participants received written and oral information about the study and provided written informed consent. At the end of the study, participants were compensated with 20€or partial course credit. The study was approved by the ethics committee of the Medical Faculty Mannheim, University of Heidelberg (ID 2022-646; approval date January 19, 2023) and preregistered at aspredicted.org (#120910, https://aspredicted.org/49kz-s6nt.pdf) before the start of data acquisition. All research was conducted according to the principles of the Declaration of Helsinki.

### 2.2 Paradigm

For the paradigm, we created 16 CT propositions, and for each proposition, 5 confirming (alternative facts) and 5 contradicting (official facts) pieces of information. For example, two of the propositions were “*Die US-Regierung hat den Angriff auf Pearl Harbor bewusst geschehen lassen*.*” (The US government deliberately allowed Pearl Harbor to happen*.) or “*Pharmakonzerne halten wirksame Medikamente zurück um weiterhin Geld zu verdienen” (Pharmaceutical companies hold back effective drugs to keep making money)*. In 16 trials, participants were first shown a CT proposition for 6 s and then rated their belief on a visual analog scale from 0 – 100 % with markers ‘*definitely not true’* (at 0%), ‘*50/50’* (at 50%), and ‘*definitely true’* (at 100%) by shifting a slider with a button box. After the rating, they were shown 5 pieces of information (either “alternative” or “official” facts for half of the propositions counterbalanced across participants) and a second presentation of the same statement followed again by the belief rating. A fixation cross was shown before each trial.

As preregistered, we calculated certainty scores *c* from graded belief ratings *b* by

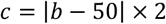

Irrespective of the direction of the truth rating, certainty c is 0 when belief was rated at 50% (‘*50/50’*) and increases towards c=100 the further the rating approaches the ends of the belief scale. Uncertainty is conceptualized as the negative (inverse) association with the certainty rating, thus, being highest at 50% and decreasing towards the ends of the belief scale.

### 2.3 MRI Scanning

MRI data was acquired at a 3T Siemens MAGNETOM PRISMAfit scanner (Siemens Healthineers, Erlangen, Germany) at the Central Institute of Mental Health in Mannheim, Germany, in a single session that included an anatomical MPRAGE scan, a 6 minute resting state scan, and the experiment. The MPRAGE was acquired with a time of repetition TR=1.3 s, echo time TE=3.03 ms, flip angle 9°, and isotropic resolution of 1×1×1 mm. Functional images were acquired with an echo-planar imaging (EPI) sequence with TR=1.02 s, TE=30 s, 63° flip angle, simultaneous multi-slice (SMS) factor 2 and GRAPPA factor 2 in 30 slices with a thickness of 3mm and 1 mm gap and 3×3 mm in-plane resolution. The experiment was self-paced, and scanning was stopped manually after completion. The average scanning duration was 19:45 min (SD: 1:03 min; range: 18:18 – 23:02 min). During scanning, respiration and pulse were monitored with built-in equipment and saved.

### 2.4 MRI Data Analysis

MRI data analysis was conducted with SPM12 (v7738) in MATLAB R2020a. The anatomical image was segmented and normalized to MNI ICBM152 non-linear asymmetric 2009b space. Functional images were slice-timing corrected, realigned, coregistered to the anatomical image and normalized to MNI space by applying the forward deformation field, rescaled to a resolution of 3×3×3 mm, and smoothed with a Gaussian kernel with FWHM=8×8×8 mm.

First level analysis was conducted with a general linear model (GLM) including condition regressors for initial CT statement presentation, its parametric modulation with the estimated certainty scores, repeated CT statements, official facts (blockwise over all 5 information presentations), alternative facts (blockwise over all 5 information presentations), and the rating phase convolved with the canonical hemodynamic response function (cHRF). Furthermore, regressors were included for events of no interest (fixation cross, button presses) convolved with the cHRF, 6 motion regressors, dummy regressors marking volumes with excessive head motion (estimated with the ART toolbox; parameters framewise displacement FD>0.5 mm, global intensity change z>4), white matter and cerebrospinal fluid signals, and physiological nuisance regressors estimated with the TAPAS PhysiO toolbox [22].

A positive contrast was put on the parametric modulation of the initial statement presentation by subsequent certainty ratings. Second level imaging analysis were conducted on the obtained contrast images with one-sample t-tests with sex and age as covariates. Uncertainty effects were estimated by a negative contrast (−1). The preregistered statistical threshold for whole-brain analysis was cluster-level significance of p <.05 with a cluster-defining threshold of p <.001 uncorrected. The effect size Hedges’ g was calculated for the imaging results based on the procedures described in Gerchen et al. [23].

To compare the identified effects with uncertainty in binary belief we plot the overlap with the respective results of Gerchen et al. [2] (their Figure 5) at the same statistical threshold (see Figure 1).

**Figure 1.**
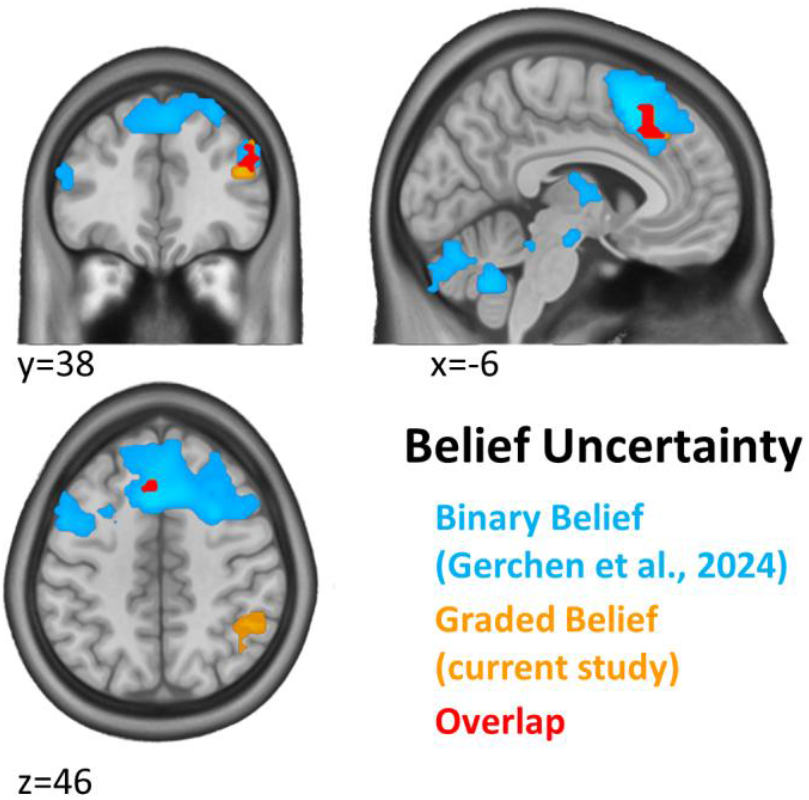
Overlap of graded and binary belief uncertainty. Overlap (red) of brain activation during initial proposition presentation associated with uncertainty estimated from graded belief ratings in the current study (orange) and brain activation during proposition presentation associated with uncertainty ratings for binary belief decisions reported in Gerchen et al. [2] (blue). Significance threshold in both analyses p=0.05 cluster-level corrected with a cluster-defining threshold (CDT) of p=0.001 unc.

## 3. Results

Participants rated their belief in the initial statement presentation at *M* = 12.68%, *SD*: 9.76% (total range 0-84%) over all 16 CT statements. A positive second level contrast on the parametric modulation of the initial statement presentation by the estimated belief certainty score showed no significant effect. For the negative contrast (uncertainty), significant clusters were found in occipital cortex, right posterior parietal cortex, right dorsolateral prefrontal cortex, and dorsomedial prefrontal cortex (Figure 1 and Table 1). When plotted on the effects associated with uncertainty ratings for binary belief decisions reported in Gerchen et al. [2], overlapping clusters are found in the dmPFC and the right dorsolateral prefrontal cortex (dlPFC, see Figure 1).

**Table 1.** Effects of parametric modulation of statement presentation with uncertainty. fMRI results at a significance threshold of p=0.05 cluster-level corrected with a cluster-defining threshold (CDT) of p=0.001 unc.

| **cluster-level** |  | **peak-level** |  |  |  |  |
| --- | --- | --- | --- | --- | --- | --- |
| **pFWE** | **k<sub>E</sub>** | **pFWE** | **t** | **Hedges' g** | **MNI (mm; x, y, z)** | **label** |
| 0.032 | 63 | 0.055 | 6.19 | 1.12 | 21, -103, 8 | occipital cortex |
| 0.001 | 127 | 0.294<br>0.999 | 5.34<br>3.86 | 0.96<br>0.70 | 48, -46, 53<br>39, -64, 53 | posterior parietal cortex |
| 0.045 | 58 | 0.488<br>0.993 | 5.02<br>4.00 | 0.91<br>0.72 | 39, 35, 20<br>48, 41, 29 | dorsolateral prefrontal cortex |
| 0.048 | 57 | 0.995<br>1.000<br>1.000 | 3.98<br>3.78<br>3.76 | 0.72<br>0.68<br>0.68 | -9, 23, 38<br>-6, 26, 50<br>-3, 29, 38 | dorsomedial prefrontal cortex |

## 4. Discussion

We identified a neural signature of belief uncertainty estimated from graded belief that overlaps in the dmPFC and right dlPFC with a signature based on the explicit rating of uncertainty about binary belief decisions. This links self-reported graded beliefs with uncertainty on the basis of brain processes, which is a relevant insight because equivalence between graded belief and confidence had previously been assumed without further evidence in the philosophical literature [16].

While identification of overlapping brain activations with fMRI cannot be taken as definite proof for the equivalence of underlying cognitive processes [24], we think that the robustness of the association provides as much empirical evidence as is possible to derive from noninvasive human brain mapping. After Gerchen et al. [2] and Bruns et al. [20], the reported results constitute our third conceptual replication of the original dmPFC belief uncertainty association of Harris et al. [19] with a modified experimental procedure. In their study, Harris et al. [19] presented propositions from different domains (for example factual, autobiographical, religious) and let participants judge by pressing one of three buttons whether they considered them true, false, or undecidable. With this categorical procedure, they identified an association with uncertainty in the dmPFC for the contrasts uncertainty > belief and uncertainty > disbelief. In an adapted version of Harris’ task, we presented propositions from the domains facts, politics, religion, CT, and superstition, and let participants rate their belief in [2]. In contrast to Harris et al. [19], in that study we separated proposition presentation, categorical true/false belief decision, and certainty judgement in this decision on a 0-100% scale. By a parametric modulation analysis we then identified a network centered at the dmPFC negatively associated during proposition presentation with subsequent certainty judgements. Thus, using a substantially different procedure, we conceptually replicated Harris’ uncertainty effect. Using a similar approach, we further replicated the dmPFC uncertainty effect for self-referential belief [20].

The identification of this association for categorical uncertainty choices [19], explicit ratings of (un)certainty in a binary belief decision [2,20], and uncertainty estimated from graded belief ratings (this study) emphasizes its robustness and might hint to a fundamental role of the underlying brain process in human belief.

While the propositions in the current study came only from the CT domain, this correspondence further suggests that the reported effects are likely domain-general rather than domain-specific (for specific CT effects see [2] and [25]).

In Gerchen et al. [2] we related our findings to the discussion of belief by Alexander Bain in the 19^th^ century and noted that the dmPFC uncertainty association identifies a neural substrate for his famous notion that “the real opposite of belief as a state of mind is not disbelief, but doubt, uncertainty” [26, p. 574].

Here we would like to expand further on this correspondence because, surprisingly, Bain already provided a fitting interpretation for the localization and function of this effect. Based on the idea that belief emerges from expectations about the contingent consequences of own actions, he assumed an intimate relationship between belief and action that extends into the cognitive domain, or in his own words: “The state in question, then, having its roots in voluntary action, branches far and wide into the realms of intelligence and speculation.” [26, p. 571]. This corresponds well with the localization of our findings. While we use the relatively broad label dmPFC, in an alternative terminology the effect falls into the anterior pre-supplementary motor area (pre-SMA; see [27,28]). Based on resting state analyses, Zhang et al. [28] showed that the anterior pre-SMA is connected with the prefrontal cortex and the caudate nucleus, but not with somatomotor areas, and Nachev et al. [29] concluded that the pre-SMA is involved in more complex and cognitive processing in comparison to more posterior regions of the supplementary motor complex.

Even more important is Bain’s description of uncertainty: “…when the only obstacle is uncertainty as to the choice of means, we are kept on the tenter-hooks of alternate expedients, encouraged and baffled by turns. Distracted by opposing considerations, keeping up an aim, and yet not making any progress towards it, we suffer all the acute misery so well known to accompany such situations of contradicting impulses.” [26, p. 574]. This corresponds astonishingly well to the modern literature on pre-SMA function. For example, Nachev et al. [30] reported that a unilateral pre-SMA lesion leads to a specific contralateral deficit in the ability to inhibit ongoing movement plans in the presence of response conflicts. Further, Usami et al. [31] showed that electrical stimulation of the pre-SMA lead to prolonged reaction times during conflict trials in a flanker task, and Wolpe et al. [32] demonstrated that a pre-SMA lesion is associated with lower response thresholds for action initiation and action choice. Using the even broader label dmPFC/dACC, which includes the pre-SMA, Clairis & Lopez-Persem [33] reviewed and compared the, partially conflicting, theories about the basic functions underlying its involvement in diverse cognitive tasks. These theories have, for example, emphasized conflict detection and conflict monitoring, the signaling of the need and value of cognitive control, the computation of prediction errors and of the likelihood of committing an error, the inhibition of actions with a high risk for negative outcomes, or the signaling of the value of foraging the environment instead of keeping with the ongoing action. While it remains unclear whether a single unifying theory on dmPFC/dACC function could exist, at least it is generally agreed on that its activity signals a need for adaptation [33].

This function of the dmPFC might thus be regarded as a core neurocognitive process that is flexibly integrated into diverse cognitive domains. In the context of belief it would correspond to, “psychologically considered, […] doubt and inquiry” [1, p. 284], and to comparable sub-functions in other domains. For example, it has recently been argued that the well-established involvement of the dmPFC in the social cognitive mentalizing network can be better explained by elevated task uncertainty in social contexts than by mental content [34].

A close correspondence to locally implemented neural computations and their basic nature might be the reasons for the robustness of the dmPFC belief uncertainty effect and its conceptual replicability that we find even with modest sample sizes.

In addition to the dmPFC we found an overlap with our results in Gerchen et al. [2] in the right dlPFC, which might, together with the right posterior parietal cortex only identified in this study, hint at the involvement of executive functions in belief uncertainty. Of note, also the dlPFC has been described to be involved in general uncertainty processing [35] and it is often assumed that the dmPFC signals a need for cognitive control that is then implemented by the dlPFC [33].

Of note, our study has clear limitations. The average belief in the presented CT statements was rather low in the investigated sample, which had mainly an academic background. However, the conducted parametric modulation analysis captures activation differences between presented propositions. Thus, it is not required that overall belief in the statements is high when at least some variability between items exists. Still, limited variability and the smaller number of statements might be reasons why the cluster is smaller than in Gerchen et al. [2].

Further, to estimate certainty scores from graded belief, we used a simple equation assuming a linear relationship across positive (belief) and negative (disbelief) directions. This does not necessarily have to be true, and the relationship might be more complex. It would be interesting to investigate the exact form of this relationship in future research.

Importantly, while the philosophical literature emphasizes the role of confidence in graded belief, we did not find a significant association in this direction, although Gerchen et al. [2] identified distinct brain networks positively associated with certainty ratings. While this is likely due to limited power in the current study, the clear, replicable, and robust association of belief uncertainty with dmPFC activation as the stronger effect speaks for a relevant role of doubt in human belief processes. Instead of assuming an accumulation of confidence until a categorical belief threshold is reached, this would be consistent with a model of graded belief in which a categorical belief is gradually put into question by increasing doubt.

## 5. Conclusions

It seems natural for humans to rate their beliefs categorically as well as on a graded scale and our results provide evidence that reported levels of graded belief reflect, at least to a certain extent, the degree of uncertainty in a belief object. Our findings correspond to the notion of Alexander Bain about the central role of doubt in belief and suggest that theoretical conceptualizations of graded belief should not focus solely on credence or confidence, but should also consider its inverse direction of doubt or uncertainty, which possesses a distinct neural and psychological reality.

## Declarations

### Ethical Statement

The study was approved by the ethics committee of the Medical Faculty Mannheim, University of Heidelberg (ID 2022-646). All participants provided written informed consent before participation.

### Author Contributions

**M.F.G**.: Conceptualization, Methodology, Formal analysis, Investigation, Resources, Data Curation, Writing - Original Draft, Writing - Review & Editing, Visualization, Supervision, Project administration, Funding acquisition **S.U**.: Conceptualization, Methodology, Formal analysis, Investigation, Data Curation, Writing - Original Draft, Writing - Review & Editing, Project administration **M.L**.: Conceptualization, Writing - Original Draft, Writing - Review & Editing, Funding acquisition **P.K**.: Conceptualization, Resources, Writing - Original Draft, Writing - Review & Editing, Supervision

### Data Availability

The datasets generated and analyzed during the current study are not publicly available due to data protection requirements but are available from the corresponding author on reasonable request.

